# From Plants to Patients: Mitochondrial Stress Signaling as a Systems Framework for Human Disease Vulnerability

**DOI:** 10.64898/2026.08.17.745221

**Authors:** F. Şeyma Gökdemir, Füsun Eyidoğan, Keshav K. Singh, Gökhan Burçin Kubat

## Abstract

Mitochondria integrate bioenergetic metabolism, redox control, genome maintenance, and stress signaling across all eukaryotes. Although plant and human mitochondria diverged substantially during evolution, both systems retain systems-level principles for sensing mitochondrial dysfunction and communicating stress signals to the nucleus. Here, we develop an integrative comparative in silico framework to evaluate whether plant mitochondrial stress signaling can provide a useful conceptual model for interpreting human mitochondrial disease vulnerability. Core *Arabidopsis thaliana* regulators representing alternative respiration, mitochondrial retrograde signaling, translational stress control, and genome surveillance were compared with functionally analogous human regulators involved in integrated stress response (ISR) signaling, mitochondrial DNA maintenance, and mitochondrial disease phenotypes. Domain architecture, protein-protein interaction topology, enrichment profiles, disease-gene associations, and promoter motif architecture were integrated to assess cross-kingdom convergence at the level of stress-response organization rather than direct orthologs. The plant network formed a compact AOX-NAC-centered stress module associated with respiratory flexibility and retrograde signaling, whereas the human network displayed expanded ISR and mtDNA maintenance modules enriched for mitochondrial disease associations. Promoter motif analyses further indicated lineage-specific transcription factor signatures but broadly comparable stress-responsive regulatory logic. Collectively, these results support the concept that plant mitochondrial stress systems represent simplified resilience-oriented architectures that can help generate experimentally testable hypotheses about failure points in human mitochondrial stress responses.

## 1. Introduction

Mitochondria are central hubs of cellular metabolism, redox homeostasis, and stress integration across eukaryotic life. Originating from an ancient alpha-proteobacterial endosymbiont, mitochondria retain deeply conserved bioenergetic architectures and signaling principles despite extensive evolutionary divergence between kingdoms (Gray, 2012; Roger et al., 2017; Jeong et al., 2025). In humans, mitochondrial dysfunction underlies a wide spectrum of inherited and acquired disorders, including primary mitochondrial diseases, neurodegenerative syndromes, metabolic disorders, and age-associated pathologies (Gorman et al., 2016; Wallace, 2018). However, the mechanistic complexity, tissue specificity, and limited therapeutic accessibility of human mitochondrial systems continue to constrain diagnosis and treatment.

Plants have evolved a notably resilient mitochondrial stress-management strategy. While operating a conserved electron transport chain and oxidative phosphorylation system, plant mitochondria also possess additional buffering, bypass, and genome surveillance mechanisms that are absent or differently organized in animals (Vanlerberghe, 2013; Sweetlove et al., 2010). Rather than representing mere evolutionary deviations, these features provide an experimental window into mitochondrial stress tolerance, signaling plasticity, and survival thresholds. Consequently, plant systems offer a powerful but underused framework for decoding mitochondrial stress responses relevant to human disease.

Although the mammalian mitochondrial respiratory chain is organized into dynamic respiratory supercomplexes, including Complex I–III–IV assemblies, electron flow remains largely dependent on the canonical cytochrome pathway and lacks an AOX-like alternative terminal oxidase. Therefore, genetic impairment or pharmacological inhibition of key respiratory complexes, particularly Complex IV, can disrupt electron transfer, promote electron leakage, increase reactive oxygen species (ROS) accumulation, and contribute to bioenergetic failure and cell death (Diaz, 2010; Scialò et al., 2017; Letts et al., 2016; Guo et al., 2017). Plants circumvent this fragility through the alternative oxidase (AOX) pathway, which branches at the ubiquinone pool and transfers electrons directly to oxygen without proton pumping (Vanlerberghe and McIntosh, 1997). Although energetically uncoupled from ATP synthesis, AOX functions as a safety valve that limits ROS accumulation under stress. Heterologous expression of AOX in mammalian cells and animal models has been shown to provide resistance to respiratory chain blockade and to alleviate phenotypes associated with Complex IV dysfunction (El-Khoury et al., 2013; Szibor et al., 2017).

Beyond bioenergetics, mitochondrial dysfunction triggers communication with the nucleus through retrograde signaling pathways that reprogram gene expression. In plants, mitochondrial retrograde signaling is coordinated primarily by membrane-tethered NAC transcription factors, particularly ANAC017 and NAC013, which activate stress-responsive nuclear programs upon mitochondrial perturbation (De Clercq et al., 2013; Ng et al., 2013). In mammals, a functionally analogous but molecularly distinct system is centered on activating transcription factor 4 (ATF4), a master regulator of mitochondrial stress and the integrated stress response (ISR) (Pakos-Zebrucka et al., 2016; Quiros et al., 2017). This parallel suggests that conservation occurs at the level of regulatory logic and network architecture rather than strict orthology of individual transcription factors.

Mitochondrial stress responses are also embedded within broader proteostasis control mechanisms. Mammals rely on multiple eIF2alpha kinases to activate the ISR, whereas plants predominantly utilize GCN2, offering a simplified but informative model for investigating stress thresholds and adaptive limits (Harding et al., 2003; Lageix et al., 2008; Lokdarshi et al., 2022). Chronic or dysregulated ISR activation is increasingly implicated in neurodegenerative and metabolic disease contexts (Costa-Mattioli and Walter, 2020). Understanding how plants maintain protective stress responses may therefore illuminate the transition from adaptive signaling to pathology in humans.

A further divergence lies in mitochondrial genome maintenance. Human mitochondrial DNA exhibits high mutation rates and limited repair capacity, leading to heteroplasmy, clonal expansion of deleterious variants, and progressive dysfunction during aging and disease (Stewart and Chinnery, 2015). Plants maintain exceptional mitochondrial genome stability despite large and recombinogenic mitochondrial genomes. The plant-specific mismatch DNA repair protein MutS HOMOLOG 1 (MSH1) protein contributes to mitochondrial and plastid genome surveillance and suppresses aberrant recombination (Abdelnoor et al., 2003; Xu et al., 2011). Deciphering how plants preserve organellar genome integrity may provide conceptual blueprints for future mtDNA-targeted strategies.

Here, we integrate plant mitochondrial stress signaling with evolutionary, network, regulatory, and disease-centered analyses to establish plants as a comparative framework for understanding human mitochondrial stress response and vulnerability. We combine domain annotation, STRING-based network topology, functional enrichment, disease mapping, and promoter motif analyses to test whether plant and human mitochondrial stress systems converge at the level of systems architecture. Throughout the manuscript, the terms analogous and convergent are used to indicate functional and regulatory similarity rather than direct gene-level orthologs.

### 2. Alternative oxidase as a mitochondrial bypass strategy

The organization of the mammalian mitochondrial electron transport chain renders human cells vulnerable to respiratory perturbations. Genetic defects, pharmacological inhibition, or environmental toxins targeting a single complex can precipitate bioenergetic failure, excessive ROS production, and cell death (Diaz, 2010; Wallace, 2018). This structural fragility contrasts with the respiratory flexibility observed in plant mitochondria.

Plants possess an AOX pathway that branches at the ubiquinone pool and directly transfers electrons to oxygen, bypassing Complexes III and IV (Vanlerberghe and McIntosh, 1997). Although this pathway does not contribute to proton gradient formation or ATP synthesis, AOX maintains electron flux under stress and prevents over-reduction of the respiratory chain. From an evolutionary perspective, AOX represents a trade-off between energetic efficiency and cellular survival.

Importantly, the relevance of AOX is not confined to plant biology. Heterologous expression of AOX in mammalian systems has demonstrated protective effects against respiratory chain inhibition, including reduced ROS accumulation and improved tolerance of Complex IV dysfunction (El-Khoury et al., 2013; Szibor et al., 2017). AOX should therefore be interpreted not as a replacement for oxidative phosphorylation, but as a conditional safeguard that preserves mitochondrial integrity during acute respiratory stress.

### 3. Mitochondrial retrograde signaling: the ANAC017-ATF4 analogy

Mitochondrial dysfunction is a potent signaling trigger that reshapes nuclear gene expression through retrograde signaling pathways. These pathways enable cells to sense mitochondrial perturbation and initiate adaptive transcriptional programs that restore homeostasis, rewire metabolism, and influence survival decisions (Quiros et al., 2017).

In plants, mitochondrial retrograde signaling is mediated by membrane-tethered NAC transcription factors, with ANAC017 emerging as a central regulator. Under basal conditions, ANAC017 is membrane-associated; upon mitochondrial stress, proteolytic cleavage releases its N-terminal transcriptional domain, enabling nuclear translocation and activation of stress-responsive genes (Ng et al., 2013; De Clercq et al., 2013). ANAC017 and NAC013 regulate detoxification enzymes, antioxidant systems, mitochondrial chaperones, and metabolic reprogramming factors.

In mammals, mitochondrial stress signaling converges on ATF4, which is preferentially translated following eIF2alpha phosphorylation and induces genes involved in amino acid metabolism, redox balance, proteostasis, and cell fate decisions (Harding et al., 2003; Pakos-Zebrucka et al., 2016). Although ANAC017 and ATF4 do not represent direct orthologs, both act as conditional nuclear effectors of mitochondrial stress. This conceptual analogy provides the rationale for the cross-kingdom comparisons presented below and is summarized in the integrated comparative framew rk shown in **Figure 1**.

**Figure 1.**
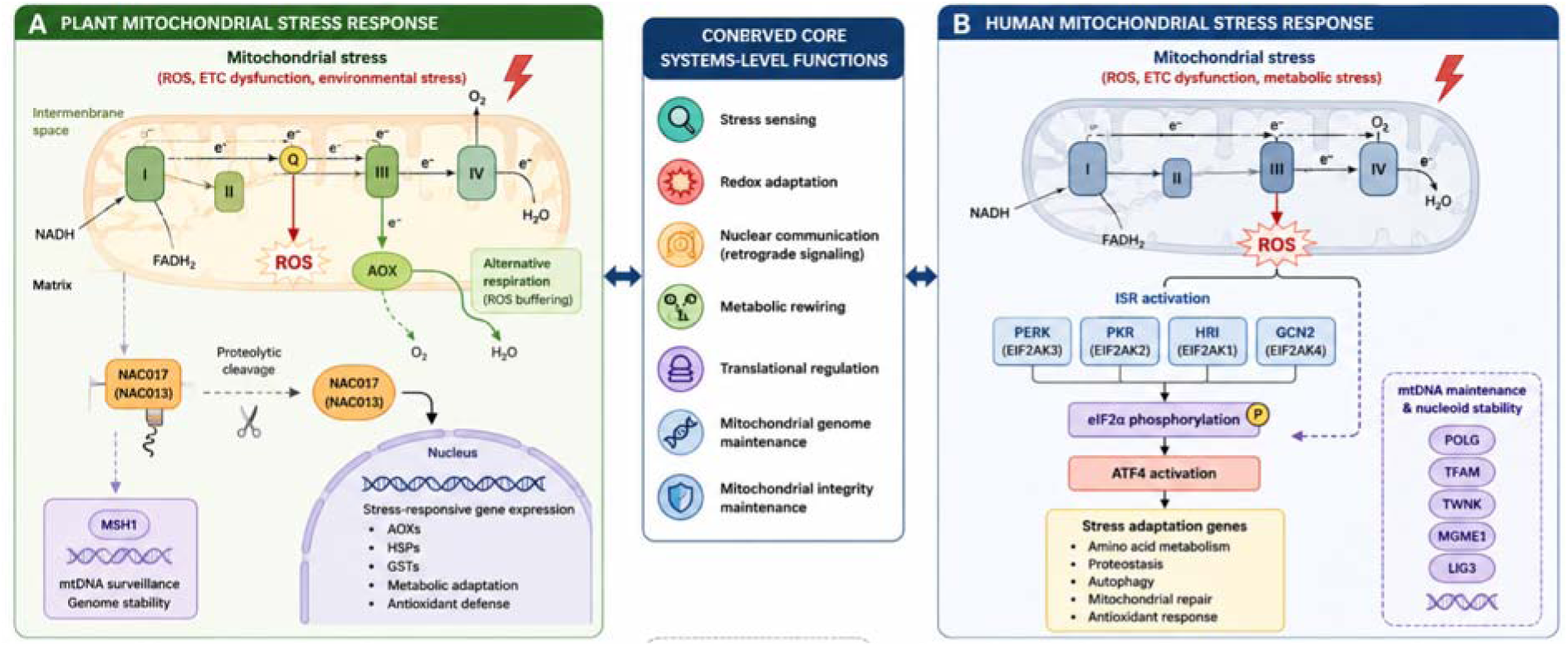
Cross-kingdom comparison of mitochondrial stress adaptation mechanisms in plants and humans. (A) Plant mitochondrial stress signaling includes AOX-mediated respiratory bypass, NAC017/NAC013-dependent retrograde signaling, and MSH1-mediated mitochondrial genome surveillance. (B) Human mitochondrial stress signaling includes ISR activation through eIF2alpha kinases, ATF4-dependent transcriptional reprogramming, and mtDNA maintenance modules involving POLG, TFAM, TWNK, MGME1, and LIG3. The central conceptual layer highlights recurrent systems-level functions, including stress sensing, redox adaptation, nuclear communication, translational regulation, metabolic rewiring, mitochondrial genome maintenance, and preservation of mitochondrial integrity. (This figure were generated using BioRender and/or artificial intelligence-assisted illustration tools).

This conceptual framework raises a central methodological question: whether plant and human mitochondrial stress systems, despite lacking broad one-to-one orthology, can be compared through shared functional layers and systems-level organization. To address this, we designed the study as an integrative comparative *in silico* analysis rather than a direct orthology search. The following analyses therefore focused on functionally analogous stress-response modules, including respiratory flexibility, mitochondria-to-nucleus communication, translational stress control, and mitochondrial genome maintenance. This design allowed us to evaluate convergence at the level of domain architecture, interaction topology, functional enrichment, disease association, and promoter regulatory logic.

## 4. Materials and Methods

### 4.1. Study design and gene selection

This study was designed as a comparative integrative *in silico* analysis rather than a strict orthology study. Genes were selected a priori to represent five biologically comparable mitochondrial stress layers: respiratory flexibility, mitochondria-to-nucleus signaling, translational stress control, mitochondrial genome maintenance, Disease and translational interpretation. For *Arabidopsis thaliana*, the core gene set included AOX1A, AOX1D, AOX2/AOX3, NAC013, NAC017, MSH1, and GCN2/RZF1-related stress components where available in STRING. For *Homo sapiens*, the core gene set included ATF4, EIF2S1, EIF2AK1, EIF2AK2, EIF2AK3, EIF2AK4, POLG, TFAM, TWNK, MGME1, LIG3, SURF1, and COX4I1. Gene selection was guided by prior experimental evidence, curated annotation, database availability, and representation of comparable mitochondrial stress-response functions rather than by one-to-one orthology (**Table 1**).

**Table 1.** Rationale for selection of core plant and human mitochondrial stress-associated genes/modules.

| Functional layer | *Arabidopsis thaliana* genes/modules | *Homo sapiens* genes/modules | Selection rationale |
| --- | --- | --- | --- |
| Respiratory flexibility and redox buffering | AOX1A, AOX1D, AOX2/AOX3 | COX4I1, SURF1 and ETC-associated vulnerability markers | Represents plant respiratory bypass capacity versus human respiratory-chain fragility and Complex IV-associated disease relevance. |
| Mitochondria-to-nucleus communication | NAC013, NAC017 | ATF4 | Captures functionally analogous retrograde/nuclear stress-response effectors without implying direct orthology. |
| Translational stress control | GCN2/RZF1-related stress components where available | EIF2S1, EIF2AK1, EIF2AK2, EIF2AK3, EIF2AK4 | Represents eIF2alpha-centered stress adaptation and ISR-like translational regulation across kingdoms. |
| Mitochondrial genome surveillance and maintenance | MSH1 | POLG, TFAM, TWNK, MGME1, LIG3 | Compares plant organellar genome surveillance with the human mtDNA replication, nucleoid stability, and repair/ligation module. |
| Disease and translational interpretation | Adaptive mitochondrial stress-resilience model | Mitochondrial disease-associated ISR-mtDNA network | Allows disease mapping on the human network while using the plant network as a resilience-oriented comparative model. |

### 4.2. Protein sequence retrieval and domain annotation

Protein sequences were retrieved from NCBI and/or UniProt entries and manually curated to retain representative protein isoforms. Comparative domain architectures were evaluated using conserved domain annotations, InterPro/Pfam domain information, and STRING functional annotation outputs. Particular attention was given to alternative oxidase ferritin-like/diiron catalytic domains, NAC DNA-binding domains, bZIP transcription factor domains, eIF2alpha kinase domains, MutS-like domains, and mitochondrial genome maintenance domains. Because plant and human systems are evolutionarily distant, the analysis emphasized functional analogy, pathway-level convergence, and systems-level regulatory logic rather than one-to-one orthology.

### 4.3. STRING-based protein-protein interaction and enrichment analysis

Protein-protein interaction networks were constructed separately for *Arabidopsis thaliana* and *Homo sapiens* using STRING (Szklarczyk et al., 2023). Networks were generated from the selected mitochondrial stress-associated gene sets using organism-specific STRING identifiers. Network topology was evaluated using node number, edge number, local clustering, and hub-like connectivity patterns derived from STRING exports and visual inspection. Functional enrichment outputs were downloaded from STRING, including Gene Ontology Biological Process, Molecular Function, Cellular Component, local network clusters, pathway terms where available, and disease-gene association terms for the human network. Enrichment terms were interpreted using false discovery rate (FDR)-corrected values provided by STRING.

### 4.4. Disease association mapping

Human disease relevance was evaluated using the STRING disease-gene association export. Enriched disease terms were filtered and interpreted based on FDR, observed gene count, and biological relevance to mitochondrial DNA maintenance, mitochondrial metabolism, neuromuscular phenotypes, and respiratory chain dysfunction. Disease enrichment results were used only for the human network because plant genes do not have direct human clinical disease annotations. The analysis was therefore framed as translational disease mapping rather than direct plant-human disease equivalence.

### 4.5. Promoter sequence extraction and motif analysis

Promoter regions were defined as 2000 bp upstream of representative transcription start sites. Arabidopsis promoter sequences were retrieved using Ensembl Plants BioMart/TAIR-compatible identifiers, whereas human promoter sequences were retrieved using Ensembl BioMart and/or UCSC-based promoter extraction when BioMart export was unstable. Duplicate sequence identifiers were removed or renamed to generate MEME-compatible FASTA files. De novo motif discovery was performed using MEME Suite (Bailey et al., 2015) with zero or one occurrence per sequence (ZOOPS), motif widths of 6-20 bp, and up to five motifs per dataset. Motif similarity searches were performed using TOMTOM (Gupta et al., 2007). Plant motifs were compared against *Arabidopsis*/plant motif databases, while human motifs were compared against human/mouse HOCOMOCO/JASPAR-compatible vertebrate motif collections (Kulakovskiy et al., 2018; Castro-Mondragon et al., 2022). TOMTOM matches were interpreted as putative transcription factor associations rather than experimentally validated binding events.

### 4.6. Integration strategy and interpretation criteria

Results from domain annotation, network topology, enrichment analysis, disease mapping, and promoter motif discovery were integrated to identify convergent systems-level themes. Evidence was considered strongest when supported by more than one analytical layer, such as network topology plus functional enrichment or promoter motif patterns plus established stress-response biology. The terms “conserved,” “analogous,” and “convergent” were used with specific meaning: conserved refers only to broad stress-response principles, analogous refers to functionally comparable but non-orthologous components, and convergent refers to similar systems-level organization arising from distinct molecular implementations.

## 5. Results

### 5.1. Comparative domain architecture of core mitochondrial stress regulators

Comparative domain architecture analysis revealed functional parallels between plant and human mitochondrial stress-associated proteins despite marked divergence at the primary sequence level. In *Arabidopsis*, AOX family proteins exhibited conserved ferritin-like diiron catalytic domains characteristic of alternative oxidases, supporting their role in ETC bypass and redox buffering. These proteins retained the canonical diiron-binding architecture associated with cyanide-resistant respiration and mitochondrial stress adaptation.

ANAC017 and NAC013 both contained conserved N-terminal NAC DNA-binding domains coupled with C-terminal membrane-associated regions, consistent with their roles as membrane-tethered transcription factors activated through stress-induced proteolytic release. This architecture supports a mechanism in which mitochondrial dysfunction triggers release of transcriptionally active NAC proteins that subsequently reprogram nuclear gene expression.

In contrast, human ATF4 possessed a canonical bZIP DNA-binding domain associated with ISR signaling. Although ANAC017/NAC013 and ATF4 are not sequence orthologs, both function as nuclear effectors of mitochondrial stress adaptation. This supports conservation at the level of regulatory logic rather than preservation of transcription factor identity.

Further divergence was observed in translational stress-control architecture. Whereas the plant system is represented primarily by GCN2-like eIF2alpha kinase signaling, the human system contains four eIF2alpha kinases: EIF2AK1/HRI, EIF2AK2/PKR, EIF2AK3/PERK, and EIF2AK4/GCN2. All four human kinases retain conserved serine/threonine kinase domains, indicating diversification of a conserved translational stress-response mechanism.

Mitochondrial genome maintenance also showed functional convergence with different molecular implementations. In plants, DNA repair protein MSH1 contains MutS-like domains associated with mitochondrial and plastid genome surveillance. In humans, mtDNA maintenance is distributed across multiple proteins, including POLG, TWNK, TFAM, MGME1, and LIG3. These findings indicate that both kingdoms maintain dedicated mitochondrial genome protection systems, although through structurally distinct molecular strategies.

### 5.2. Comparative network topology of plant and human mitochondrial stress signaling

STRING-based PPI analysis revealed clear topological differences between the plant and human mitochondrial stress-associated networks. The *Arabidopsis* core network consisted of seven proteins connected by four high-confidence interaction edges, forming a compact architecture centered around AOX- and NAC-associated signaling components. AOX1A, AOX3/AOX-associated nodes, NAC013, and NAC017 represented the principal connected module, suggesting a functional coupling between respiratory flexibility and mitochondrial retrograde signaling. MSH1 appeared more peripheral, consistent with a genome surveillance function that is related to, but partly separable from, the AOX-NAC stress module.

The human network contained thirteen proteins connected by twenty-nine interaction edges, indicating substantially greater connectivity and modularity. Two major modules were apparent: an ISR module involving ATF4, EIF2S1, and EIF2AK1-4, and an mtDNA maintenance module involving TFAM, POLG, TWNK, MGME1, and LIG3. TFAM and POLG emerged as prominent hub-like proteins within the mtDNA maintenance module, whereas ATF4 and eIF2alpha kinase-associated interactions formed the ISR signaling axis.

These topological differences indicate that plant mitochondrial stress signaling is organized as a compact adaptive architecture, whereas the human system integrates mitochondrial dysfunction into a broader multilayered stress network. Despite differences in individual proteins, both networks contained components associated with stress sensing, redox adaptation, nuclear communication, translational regulation, and mitochondrial integrity maintenance (**Figure 2**).

**Figure 2.**
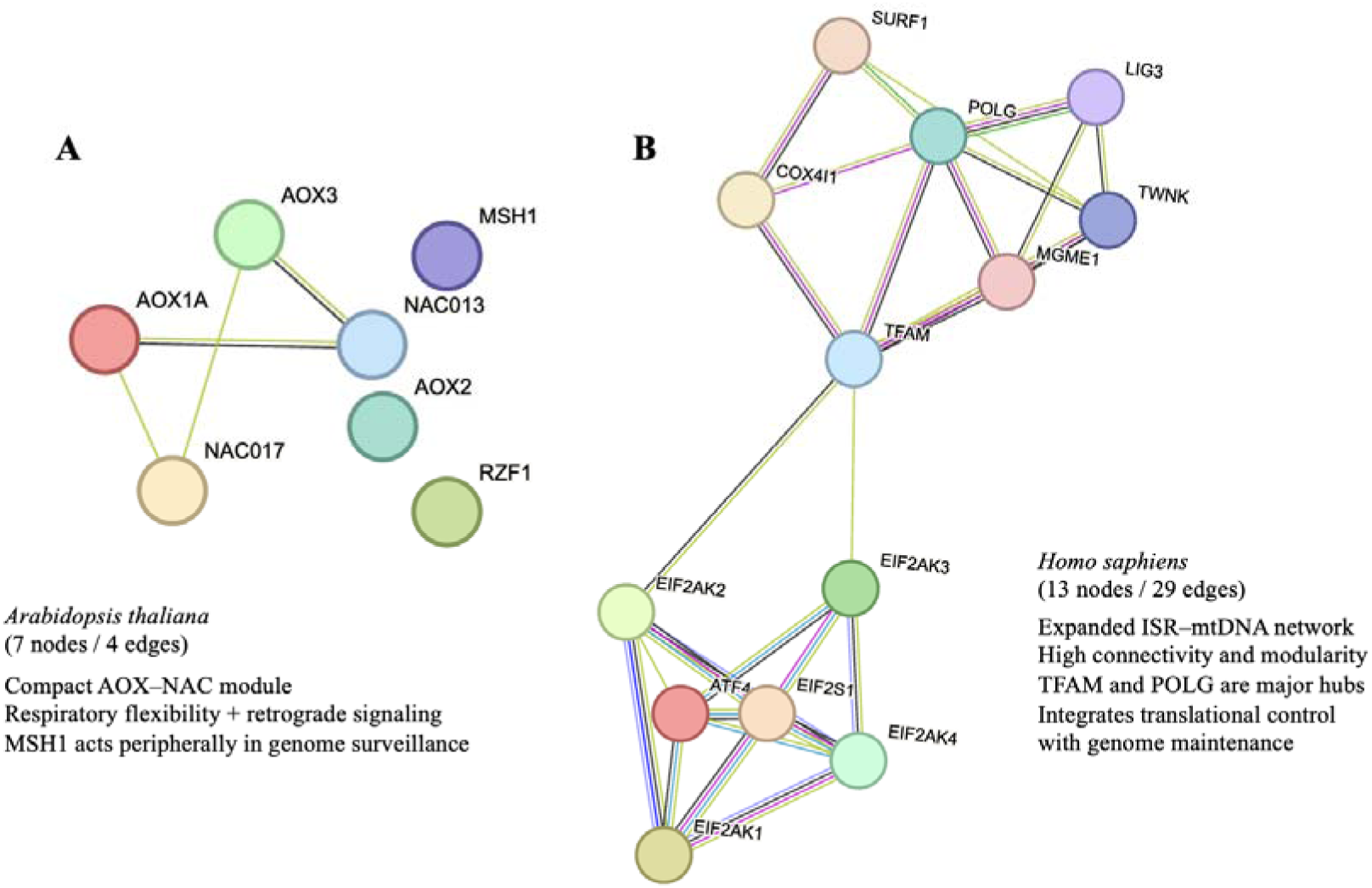
Comparative protein-protein interaction topology of mitochondrial stress-associated regulators in plants and humans. (A) STRING-based PPI network constructed using core Arabidopsis thaliana mitochondrial stress-associated proteins. The plant network exhibits a compact AOX-NAC-centered architecture associated with respiratory flexibility and retrograde signaling. (B) STRING-based PPI network constructed using core Homo sapiens mitochondrial stress-associated proteins. The human network shows greater connectivity and modularity, dominated by ISR-associated kinases and mtDNA maintenance proteins. TFAM and POLG form major hub-like components of the mtDNA maintenance module, while ATF4 and eIF2alpha kinases form the ISR signaling axis.

### 5.3. Comparative enrichment and system-level network analysis

Comparative enrichment analyses revealed substantial differences in the organizational complexity of plant and human mitochondrial stress-associated regulatory networks. The *Arabidopsis* netw rk exhibited a compact and specialized architecture primarily associated with AOX-mediated respiratory flexibility and NAC-dependent retrograde signaling. In contrast, the human network displayed a highly interconnected framework integrating translational regulation, ISR signaling, and mitochondrial genome maintenance (**Table 2**).

**Table 2.** Comparative enrichment analysis of plant and human mitochondrial stress-associated networks.

| Category | *Arabidopsis thaliana* network | *Homo sapiens* network | Biological interpretation |
| --- | --- | --- | --- |
| Core stress architecture | Compact AOX-NAC module | Expanded ISR-mtDNA maintenance network | Plants utilize a specialized adaptive stress module, whereas humans display a highly interconnected stress-regulatory architecture. |
| Enriched biological processes | Alternative respiration, oxidative stress response, retrograde signaling | Translational regulation, ISR activation, mitochondrial stress adaptation | Plant systems prioritize respiratory flexibility, whereas human systems integrate mitochondrial dysfunction with translational control. |
| Major hub components | AOX1A, NAC013/NAC017 | TFAM, POLG, EIF2AK3, ATF4 | Human mitochondrial stress networks are more centralized and modular. |
| Dominant molecular functions | Oxidoreductase activity, ubiquinol oxidase activity | Kinase activity, stress-induced transcriptional regulation | Indicates divergence between metabolic buffering and signaling amplification. |
| Conserved signaling themes | ROS sensing, nuclear communication, stress adaptation | ROS sensing, ISR signaling, nuclear stress communication | Suggests conservation of mitochondria-to-nucleus signaling logic. |
| Genome maintenance components | MSH1-mediated surveillance | POLG-TFAM-TWINK-MGME1 axis | Human mtDNA integrity depends on a broader dedicated maintenance system. |
| Disease/translational association | Indirect stress adaptation model | Direct association with mitochondrial diseases and neurodegeneration | Human modules show strong disease-linked enrichment. |
| Network topology | Low connectivity, compact organization | High connectivity, modular architecture | Supports divergence between adaptive and pathological stress-response architectures. |

In *Arabidopsis*, enrichment terms were associated with alternative respiration, oxidoreductase activity, oxidative stress adaptation, and mitochondrial retrograde communication. Domain enrichment further highlighted ferritin-like and diiron-binding catalytic features within AOX proteins, supporting their role as redox-regulatory components.

By contrast, the human network showed broader enrichment for translational regulation, eIF2alpha phosphorylation, ISR activation, stress-induced transcriptional adaptation, mitochondrial organization, and mtDNA maintenance. KEGG and Reactome-associated terms further linked the human network to mitophagy, neurodegeneration-related processes, metabolic stress, and ISR-mediated signaling cascades. This enrichment pattern is consistent with a disease-relevant mitochondrial stress network in which mitochondrial integrity and nuclear stress signaling are highly coupled.

### 5.4. Disease-associated convergence of the human mitochondrial stress network

Disease-gene association analysis demonstrated that the human mitochondrial stress-associated network is strongly enriched for primary mitochondrial disease phenotypes, particularly disorders linked to mtDNA maintenance and replication failure. The most significant disease associations included mitochondrial DNA depletion syndrome and chronic progressive external ophthalmoplegia, both associated with TWNK, MGME1, and POLG. These findings indicate that the human core network directly overlaps with clinically relevant mitochondrial disease mechanisms rather than representing a generic stress-response module (**Table 3**).

**Table 3.**
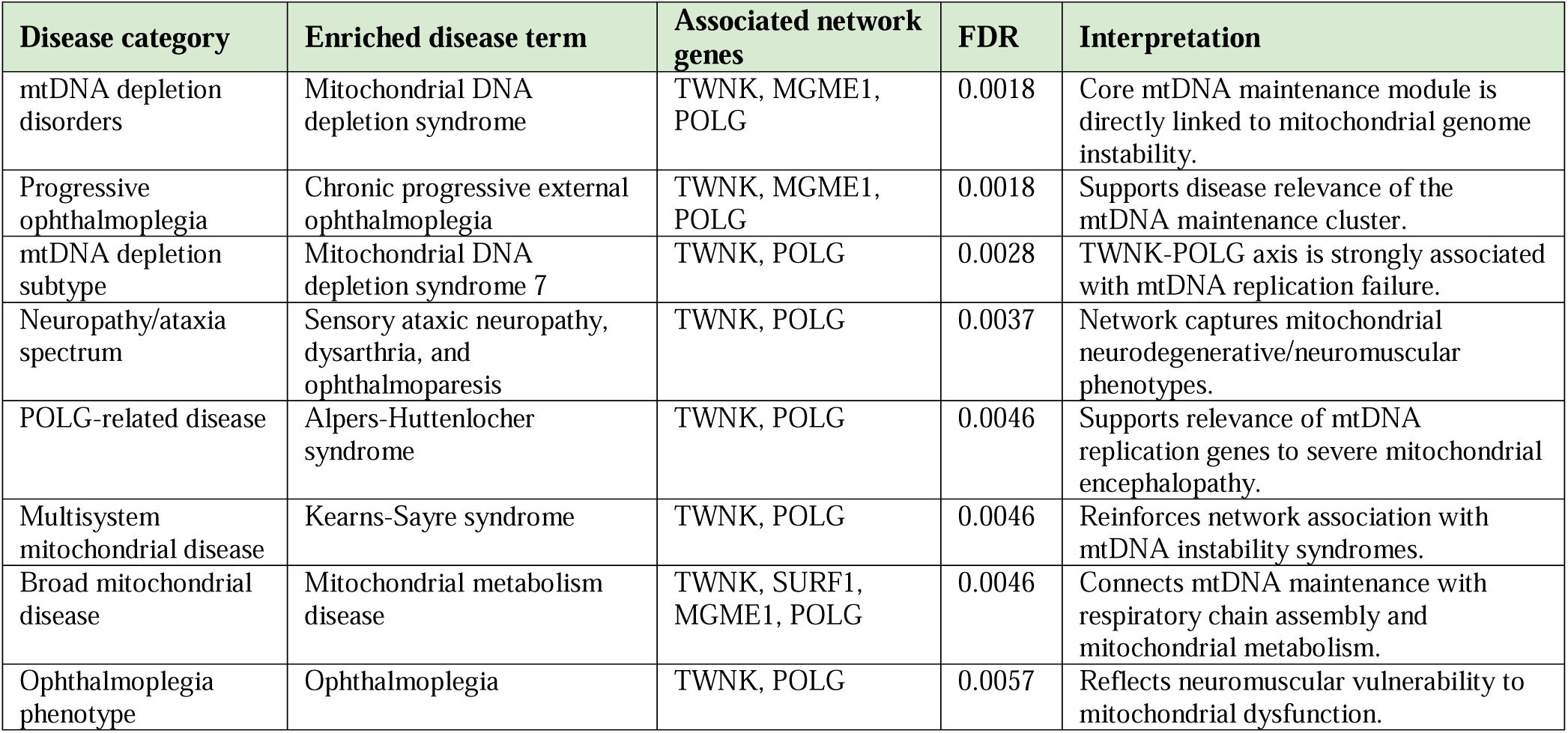
Disease relevance mapping of the human mitochondrial stress-associated network.

Several enriched disease terms were centered on the TWNK-POLG axis, including mitochondrial DNA depletion syndrome 7, sensory ataxic neuropathy, dysarthria and ophthalmoparesis, Alpers-Hutten Locher syndrome, Kearns-Sayre syndrome, and ophthalmoplegia. This pattern emphasizes the central role of mitochondrial DNA replication and genome stability in human mitochondrial pathology. In parallel, enrichment of mitochondrial metabolism disease involving TWNK, SURF1, MGME1, and POLG links genome maintenance defects with respiratory chain dysfunction and mitochondrial metabolic impairment.

These disease-associated enrichments distinguish the human network from the plant network. Whereas the plant mitochondrial stress module appears organized around adaptive resilience mechanisms such as AOX-mediated respiratory flexibility and NAC-dependent retrograde signaling, the human network contains disease-enriched modules associated with mtDNA instability, respiratory dysfunction, and neuromuscular vulnerability.

### 5.5. Cross-kingdom functional convergence of mitochondrial stress architectures

Comparative pathway and domain analyses revealed that plant and human mitochondrial stress-associated systems exhibit functional convergence despite major differences in their molecular components. In *Arabidopsis*, InterPro and domain enrichment highlighted AOX-associated ferritin-like and diiron catalytic modules shared among AOX1A, AOX2/AOX3, and AOX1D. These findings reinforce the role of AOX proteins as central adaptive components mediating respiratory flexibility and oxidative stress buffering (**Table 4**).

**Table 4.**
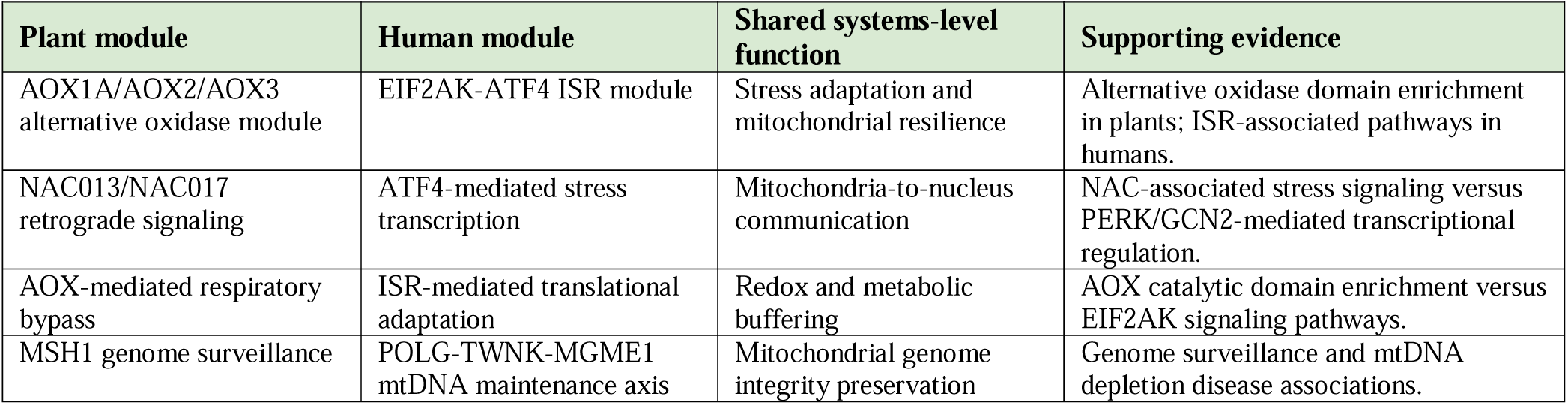
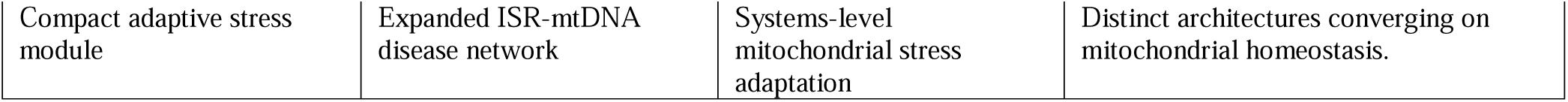
Cross-kingdom functional convergence of mitochondrial stress adaptation systems.

In the human network, Reactome and pathway enrichment emphasized cellular stress responses, eIF2alpha kinase signaling, PERK-dependent transcriptional regulation, EIF2AK4/GCN2-mediated amino acid stress signaling, and mtDNA maintenance. Core ISR-associated proteins, including EIF2AK1, EIF2AK3, EIF2AK4, EIF2S1, and ATF4, formed interconnected signaling modules linked to translational adaptation and stress-responsive gene expression.

Importantly, AOX proteins and ISR kinases are not orthologous. Nevertheless, both systems fulfill analogous adaptive functions during mitochondrial dysfunction. The plant AOX pathway buffers electron overflow and limits ROS accumulation through respiratory bypass, whereas the human ISR coordinates translational attenuation and stress-responsive transcriptional adaptation. This pattern suggests convergence at the level of systems architecture rather than direct sequence conservation.

A similar pattern was observed for mitochondrial genome maintenance. In plants, MSH1 represents a specialized organellar genome surveillance component associated with mitochondrial stability and recombination control. In humans, genome integrity functions are distributed across a broader mtDNA maintenance module including POLG, TWNK, MGME1, TFAM, and LIG3. Disease enrichment further demonstrated that this human module is strongly associated with mitochondrial depletion syndromes and neuromuscular mitochondrial disorders. The integrated systems-level model summarizing the convergence between plant adaptive mitochondrial stress modules and human ISR– mtDNA maintenance networks is presented in **Figure 3**.

**Figure 3.**
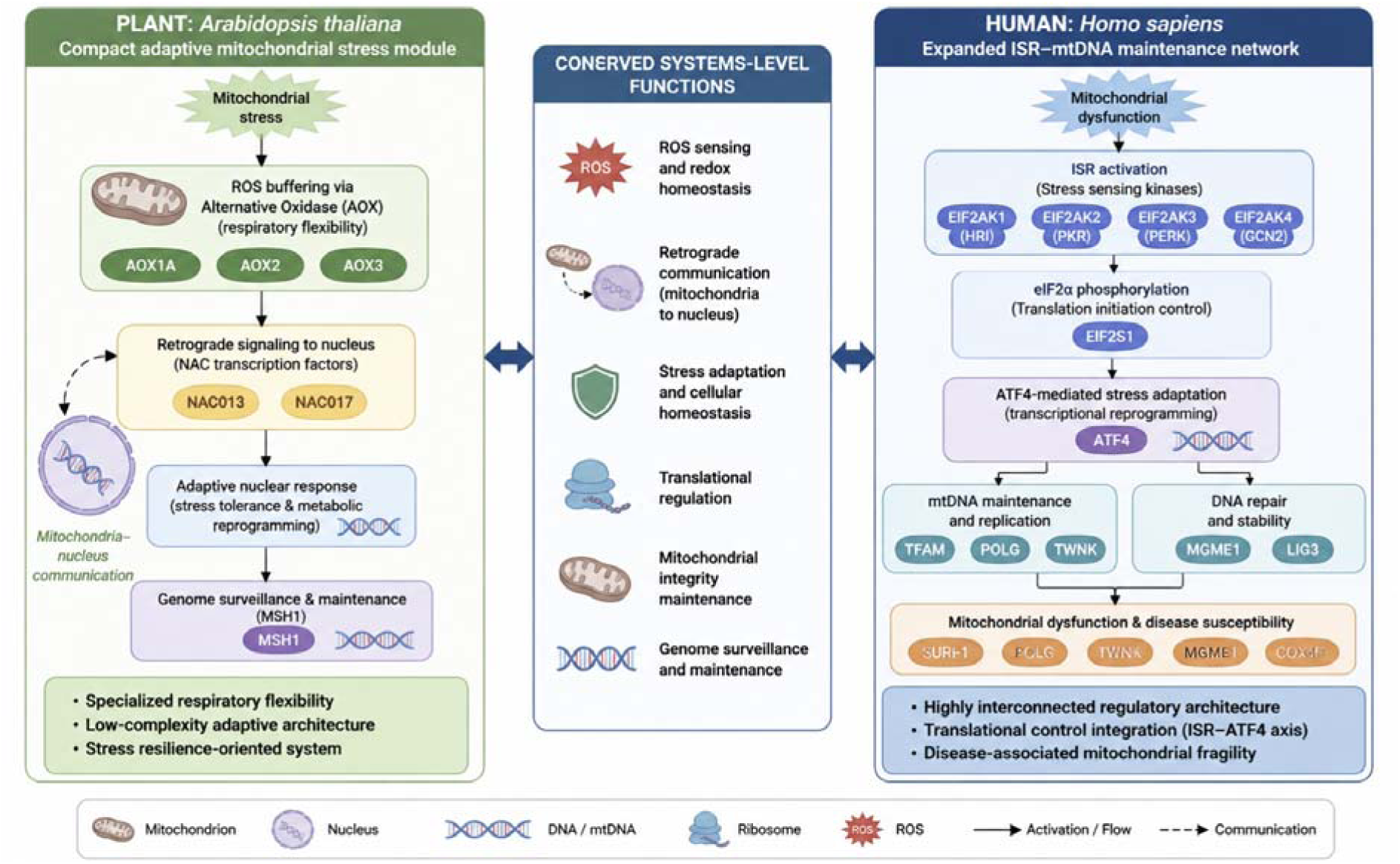
Integrated system-level model of cross-kingdom mitochondrial stress adaptation and disease-associated vulnerability. (This figure were generated using BioRender and/or artificial intelligence-assisted illustration tools).

### 5.6. Comparative promoter motif analyses suggest lineage-specific but broadly stress-responsive regulatory architectures

MEME-based promoter motif analysis identified enriched cis-regulatory motifs in both *Arabidopsis* and human mitochondrial stress-associated gene sets. In *Arabidopsis*, the three representative motifs displayed strong enrichment, with E-values ranging from 2.1 x 10^-33 to 5.8 x 10^-26 and motif occurrences distributed across 16-20 promoter sites. These motifs were selected based on statistical significance, site distribution, and biological plausibility in relation to mitochondrial stress signaling. TOMTOM-based motif comparison suggested putative similarity to stress-associated transcription factor families, including ERF/AP2-, NAC-, C2H2-, and DOF-related regulatory signatures (**Table 5**).

**Table 5.** Comparative regulatory themes identified across plant and human mitochondrial stress-associated promoter architectures.

| Regulatory layer | *Arabidopsis* | Human | Shared systems-level implication |
| --- | --- | --- | --- |
| Stress-responsive TF families | NAC, ERF/AP2 | FOX, ZNF | Transcriptional adaptation. |
| Mitochondria-to-nucleus signaling | ANAC017/NAC013 | ISR-associated signaling | Retrograde communication. |
| Oxidative stress adaptation | AOX-associated pathways | Stress-responsive transcription | Redox homeostasis. |
| Regulatory plasticity | C2H2/DOF motifs | PRDM/ZNF motifs | Transcriptional rewiring. |
| Mitochondrial genome integrity | MSH1 surveillance | POLG/TWINK maintenance | Genome stability. |

In the human promoter dataset, motif enrichment was more pronounced, with representative motifs showing E-values ranging from 1.2 x 10^-121 to 1.9 x 10^-88 and broader site representation across the analyzed sequences. TOMTOM-based annotation suggested putative associations with FOX-, zinc finger (ZNF)-, and PRDM-related transcriptional regulatory signatures. These transcription factor families are linked to stress-responsive transcriptional rewiring, genome regulatory control, and adaptive nuclear responses to cellular dysfunction (**Figure 4**).

**Figure 4.**
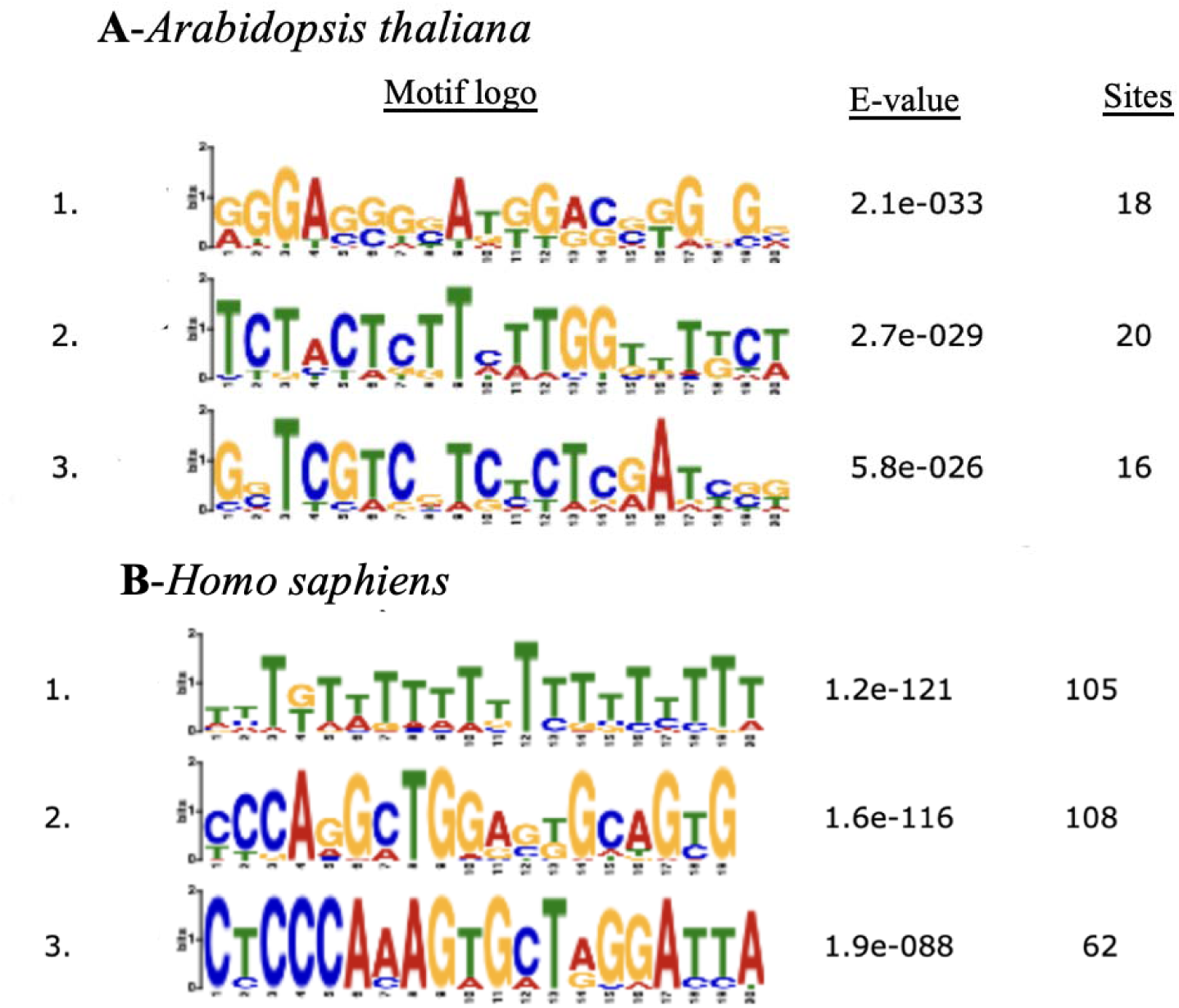
Comparative promoter motif architecture of plant and human mitochondrial stress-associated genes. Representative enriched cis-regulatory motifs identified by MEME analysis in upstream promoter regions of *Arabidopsis thaliana* and *Homo sapiens* mitochondrial stress-associated genes. (A) The *Arabidopsis* promoter set showed three enriched motifs with E-values ranging from 2.1 x 10^-33 to 5.8 x 10^-26. TOMTOM-based motif comparison suggested putative similarity to stress-associated plant transcription factor families, including ERF/AP2-, NAC-, C2H2-, and DOF-related regulatory signatures. (B) The human promoter set showed stronger motif enrichment, with E-values ranging from 1.2 x 10^-121 to 1.9 x 10^-88 and broader motif distribution across the analyzed promoters. Comparative annotation suggested associations with FOX-, ZNF-, and PRDM-related transcriptional regulatory signatures. These results indicate lineage-spec fic promoter motif compositions with convergence on stress-responsive nuclear transcriptional regulation.

Although the identified motif families differed between *Arabidopsis* and human datasets, both systems showed enrichment of promoter architectures plausibly associated with stress-responsive nuclear regulation. Therefore, plant and human mitochondrial stress pathways should not be interpreted as conserving identical transcription factors or cis-regulatory elements. Instead, the results support convergence at the level of regulatory logic: both systems coordinate mitochondrial dysfunction with nuclear transcriptional reprogramming, redox adaptation, and mitochondrial integrity maintenance.

These promoter motifs complement the PPI network, enrichment, and disease-mapping analyses by adding a hypothesis-generating cis-regulatory layer to the cross-kingdom comparison. Because motif discovery and TOMTOM similarity searches do not confirm transcription factor binding, these results should be interpreted as putative regulatory signatures that require future validation by promoter-reporter assays, ChIP-based datasets, or perturbation experiments.

## 6. Discussion

Mitochondria are increasingly recognized not only as bioenergetic organelles but also as central regulators of cellular stress adaptation, metabolic plasticity, redox homeostasis, and inter-organellar communication. Recent advances in mitochondrial systems biology have demonstrated that mitochondrial dysfunction activates highly coordinated retrograde signaling pathways capable of reshaping nuclear transcriptional programs, translational control, proteostasis, and genome maintenance mechanisms across eukaryotic lineages (Quirós et al., 2017; Pakos-Zebrucka et al., 2016; Ryoo, 2024). In the present study, comparative systems-level analyses revealed that *Arabidopsis thaliana* and *Homo sapiens* possess evolutionary distinct yet functionally convergent mitochondrial stress-response architectures. Although the underlying molecular components differ substantially between kingdoms, both systems appear to preserve conserved adaptive principles linking mitochondrial dysfunction to nuclear stress adaptation.

One of the most notable findings of this study was the marked architectural distinction between the compact AOX-centered adaptive module identified in plants and the highly interconnected ISR– mtDNA maintenance network observed in humans. In Arabidopsis, the mitochondrial stress network was organized around alternative oxidases (AOX1A/AOX3), NAC transcription factors (ANAC013/ANAC017), and the genome surveillance regulator MSH1. This relatively low-complexity architecture contrasted sharply with the human network, where ISR-associated kinases (EIF2AK1–4), ATF4-mediated transcriptional regulation, and mtDNA maintenance factors including TFAM, POLG, TWNK, MGME1, and LIG3 formed densely connected regulatory modules. These observations support the emerging concept that conserved stress-adaptive logic may persist across evolution despite divergence in gene identity and regulatory composition (Van Aken and Pogson, 2017).

The plant AOX-centered network identified in this study highlights the importance of respiratory flexibility as a core mitochondrial resilience mechanism. AOX proteins provide an alternative electron transport route that bypasses complexes III and IV of the classical mitochondrial electron transport chain, thereby preventing excessive electron over-reduction and limiting ROS accumulation under stress conditions (Vanlerberghe, 2013; Selinski et al., 2018). Previous studies demonstrated that AOX activity contributes to drought tolerance, metabolic adjustment, thermogenesis, and maintenance of photosynthetic efficiency during abiotic stress (Del-Saz et al., 2018; Moore et al., 2013). Importantly, AOX-mediated buffering may represent a unique evolutionary solution enabling plants to tolerate prolonged environmental fluctuations while maintaining mitochondrial integrity and redox balance. The prominent positioning of AOX1A and AOX3 within the present network analyses further supports the hypothesis that respiratory bypass systems function as major adaptive hubs within plant mitochondrial stress biology.

In contrast, mammalian systems appear to rely more heavily on integrated translational control mechanisms rather than respiratory bypass flexibility. Our analyses identified EIF2AK family kinases, EIF2S1, and ATF4 as central hubs linking mitochondrial dysfunction to adaptive nuclear responses. This observation is highly consistent with recent studies describing the mitochondrial integrated stress response (ISRmt) as a major adaptive signaling axis activated by oxidative phosphorylation defects, proteotoxic stress, mitochondrial translation imbalance, and ROS accumulation (Quirós et al., 2017; Fessler et al., 2020; Guo et al., 2020; Ryoo, 2024). Phosphorylation of eIF2α by stress-sensing kinases suppresses global protein translation while selectively promoting ATF4-dependent adaptive transcriptional programs controlling amino acid metabolism, antioxidant defense, autophagy, and proteostasis (Pakos-Zebrucka et al., 2016). Recent evidence further demonstrated that chronic ISR activation contributes directly to mitochondrial disease progression and tissue dysfunction, particularly in metabolically demanding tissues such as muscle and nervous systems (Jeong et al., 2025).

A particularly important conceptual outcome of this study is the systems-level convergence observed between plant retrograde signaling and mammalian ISR-mediated mitochondrial communication. Although ANAC017 and ATF4 are not sequence orthologs, both function as master regulators coupling mitochondrial dysfunction to nuclear transcriptional adaptation. In plants, ANAC017 is activated through stress-induced proteolytic cleavage and subsequently translocates to the nucleus to regulate mitochondrial dysfunction stimulon (MDS) genes associated with oxidative stress tolerance, detoxification, and metabolic adaptation (Ng et al., 2013; De Clercq et al., 2013). Similarly, ATF4 integrates mitochondrial dysfunction with nuclear stress adaptation through ISR-dependent transcriptional remodeling (Quirós et al., 2017). This functional analogy strongly suggests that evolutionarily distant organisms may preserve conserved adaptive regulatory logic even when the molecular machinery executing these responses differs substantially. Rather than direct orthology, the observed similarities therefore represent convergence at the level of systems architecture and stress-response strategy.

Another major finding of this study was the strong enrichment of mtDNA instability-associated disease pathways within the human mitochondrial stress network. Disease ontology analyses revealed highly significant associations with mitochondrial DNA depletion syndrome, chronic progressive external ophthalmoplegia, Alpers–Huttenlocher syndrome, and broader mitochondrial metabolism disorders. Central hubs included POLG, TWNK, MGME1, and TFAM, all of which are critically involved in mtDNA replication, nucleoid stability, and mitochondrial genome maintenance. Defects in these genes compromise oxidative phosphorylation capacity, increase oxidative damage, and promote progressive neurodegenerative and neuromuscular phenotypes (Viscomi and Zeviani, 2017; 2020; Gorman et al., 2016). The clustering of these genes within highly interconnected network modules further supports the notion that mammalian mitochondrial resilience depends heavily on coordinated genome surveillance and translational adaptation systems.

Interestingly, the comparatively lower topological complexity of the plant network may itself represent an adaptive feature. Plants continuously encounter fluctuating environmental conditions including drought, salinity, temperature shifts, and oxidative stress, requiring rapid metabolic flexibility and stress buffering capacity. AOX-mediated electron overflow buffering and NAC-dependent retrograde signaling may therefore constitute an efficient resilience-oriented architecture optimized for survival under chronic environmental instability (Vanlerberghe, 2013; Van Aken and Pogson, 2017). Mammalian systems, by contrast, appear to prioritize tightly coordinated translational regulation, mitochondrial proteostasis, and genome maintenance mechanisms necessary for preserving energetic stability in specialized tissues with high ATP demand. This distinction may reflect fundamental differences in ecological strategy and organismal physiology.

The promoter motif analyses further suggested the existence of broadly comparable stress-responsive regulatory themes across kingdoms. MEME and TOMTOM analyses identified putative NAC-, AP2/ERF-, DOF-, and C2H2-associated motifs within Arabidopsis promoters, whereas human promoters displayed putative FOX-, ZNF-, PRDM-, and stress-associated vertebrate regulatory signatures. Although the specific transcription factor families differed substantially between species, both promoter datasets contained motif patterns consistent with stress adaptation, transcriptional plasticity, and mitochondrial communication. These observations suggest that promoter-level convergence, if confirmed experimentally, is more likely to occur through preservation of stress-responsive regulatory logic than through direct conservation of cis-regulatory sequences.

The translational implications of these findings are particularly noteworthy. Increasing evidence suggests that controlled mitochondrial stress can activate adaptive resilience pathways, a phenomenon frequently referred to as mitohormesis (Wang and Zhang, 2025). Within this framework, plant mitochondrial systems may provide simplified biological models for investigating adaptive mitochondrial stress tolerance mechanisms relevant to human disease biology. In particular, AOX-mediated respiratory buffering has attracted growing interest as a potential therapeutic concept for reducing oxidative overload associated with ETC dysfunction (El-Khoury et al., 2013). Experimental studies introducing AOX into mammalian systems demonstrated partial rescue of respiratory defects and oxidative stress phenotypes, highlighting the translational relevance of plant-inspired mitochondrial resilience mechanisms. Recent perspective work on mitochondria transfer and transplantation further emphasizes the importance of mitochondrial delivery, integration, stability, and cellular stress resilience as emerging themes in mitochondrial medicine (Brestoff et al., 2025; Kubat et al., 2025).

Several limitations should be acknowledged. First, all analyses performed in this study were computational and based on publicly available interaction databases, enrichment analyses, promoter prediction tools, and curated literature-derived gene sets. Second, the identified relationships represent functional convergence rather than direct mechanistic equivalence or molecular orthology. Third, promoter motif enrichment analyses do not experimentally confirm transcription factor binding or regulatory activity and should be considered hypothesis-generating. Fourth, STRING-derived interaction networks may not fully capture tissue-specific, developmental, or stress-condition-dependent mitochondrial responses. Finally, the relatively small gene sets were intentionally selected to represent core stress-response layers, but broader genome-scale analyses will be required to evaluate the generality of the proposed framework. Nevertheless, systems biology approaches remain valuable for identifying conserved adaptive principles and generating experimentally testable hypotheses across evolutionary distant organisms.

Collectively, the present study demonstrates that plant and human mitochondrial stress systems exhibit substantial convergence at the level of adaptive systems architecture despite major evolutionary divergence in their molecular components. The compact AOX-NAC resilience module identified in plants and the ISR-mtDNA maintenance network identified in humans represent alternative evolutionary strategies for maintaining mitochondrial integrity under stress. These findings support the emerging view that cross-kingdom comparative mitochondrial biology may provide useful conceptual frameworks for understanding mitochondrial resilience, adaptive stress signaling, and disease-associated mitochondrial fragility.

## 7. Conclusions

Mitochondrial dysfunction is increasingly recognized as a common denominator of aging, neurodegeneration, metabolic disorders, and primary mitochondrial diseases. At the same time, recent advances in mitochondrial biology have shifted the field away from viewing mitochondria solely as bioenergetic organelles toward recognizing them as dynamic signaling hubs that coordinate cellular adaptation, stress perception, and survival decisions (Quirós et al., 2017; Costa-Mattioli & Walter, 2020; Kubat et al., 2025). Within this emerging framework, understanding how organisms maintain mitochondrial function under stress has become as important as understanding how mitochondrial dysfunction arises.

By integrating protein interaction networks, functional enrichment analyses, disease associations, and promoter-level regulatory architectures, this study demonstrates that plant and human mitochondrial stress systems converge at the level of adaptive design despite substantial divergence in their molecular components. In plants, the AOX–NAC–MSH1 module represents a resilience-oriented architecture centered on respiratory flexibility, retrograde signaling, and mitochondrial genome surveillance. In humans, the ISR–ATF4–POLG/TWNK axis coordinates translational adaptation, mitochondrial genome maintenance, and disease-associated stress responses. Although these systems rely on different molecular machinery, both ultimately serve the same biological objective: preserving mitochondrial integrity in the face of environmental, metabolic, or genetic perturbations.

Recent studies have highlighted the central role of mitochondrial retrograde signaling in coordinating organelle-to-nucleus communication across eukaryotes (De Clercq et al., 2013; Ng et al., 2013; Van Aken & Pogson, 2017; Barreto et al., 2022). Likewise, growing evidence suggests that mitochondrial resilience depends not only on energy production but also on the ability to dynamically buffer oxidative stress, reprogram nuclear gene expression, and maintain organelle homeostasis (Vanlerberghe, 2013; Selinski et al., 2018). Our findings extend these concepts by suggesting that such adaptive principles may be conserved across kingdoms even when the underlying genes are not.

Importantly, the present work does not propose that plant mitochondria directly model human mitochondrial diseases. Rather, it suggests that plant mitochondrial systems may model the adaptive boundaries that prevent mitochondrial dysfunction from becoming pathological. While human mitochondrial disorders frequently emerge when stress-buffering mechanisms become overwhelmed or insufficient, plant systems have evolved highly effective resilience strategies that allow long-term survival under chronic environmental stress. Understanding these strategies may therefore reveal fundamental principles of mitochondrial robustness that are difficult to discern from disease states alone.

The increasing interest in mitohormesis, stress-induced adaptation, and mitochondrial resilience biology further strengthens the relevance of this perspective (Costa-Mattioli & Walter, 2020; Wang & Zhang, 2025). Rather than focusing exclusively on repairing damaged mitochondria, future therapeutic approaches may increasingly seek to enhance endogenous adaptive capacity, stress buffering, and organelle quality control. In this context, plant mitochondrial stress systems offer a rich source of alternative evolutionary solutions that may inspire novel concepts in mitochondrial medicine.

Although the present study is computational and hypothesis-generating in nature, the convergence observed across network topology, disease mapping, functional enrichment, and promoter architecture analyses supports the robustness of the proposed framework. Future experimental validation of AOX-mediated buffering, NAC-dependent retrograde signaling, and MSH1-associated mitochondrial genome surveillance will be necessary to test the translational relevance of these observations.

Ultimately, the most informative question in cross-kingdom mitochondrial biology may not be whether plants and humans use the same genes, but whether they solve the same biological problem. Our analysis suggests that they do. Cross-kingdom mitochondrial biology may therefore reveal not only why mitochondria fail, but also how mitochondrial failure can be buffered, delayed, and adaptively integrated into cellular survival programs. By shifting attention from molecular conservation to resilience architecture, this framework provides a new perspective for understanding mitochondrial adaptation, disease vulnerability, and the future development of mitochondria-centered therapeutic strategies.

## Data availability

All analyses were performed using publicly available protein sequences, promoter sequences, and database-derived annotations. Edited gene lists, STRING exports, enrichment tables, MEME/TOMTOM motif outputs, and figure source files are available upon request..

## Author note on interpretation

Computational motifs and network results should be interpreted as hypothesis-generating. Terms such as “putative,” “analogous,” and “convergent” are used intentionally to avoid overstatement of unvalidated regulatory interactions or direct orthology.

## CRediT Author Statement

F. Şeyma Gökdemir: Conceptualization, Data curation, Formal analysis, Investigation, Resources, Supervision, Validation, Visualization, Writing – original draft. Füsun Eyidoğan: Conceptualization, Project administration, Supervision, Validation, Writing – review & editing. Keshav K. Singh: Supervision, Validation, Writing – review & editing. Gökhan Burçin Kubat: Supervision, Validation, Writing – review & editing.

## Declaration of generative AI and AI-assisted technologies

During the preparation of this manuscript, the author used ChatGPT for language refinement, conceptual organization, and visual ideation. AI-assisted outputs were used as preliminary references during the development of schematic figures. The final artwork was manually edited, scientifically verified, and reformatted by the author using PowerPoint and BioRender. The author takes full responsibility for the accuracy, integrity, and originality of the manuscript and all figures.

## References

Abdelnoor, R. V., Yule, R., Elo, A., Christensen, A. C., Meyer-Gauen, G., & Mackenzie, S. A. (2003). Substoichiometric shifting in the plant mitochondrial genome is influenced by a gene homologous to MutS. Proceedings of the National Academy of Sciences, 100(10), 5968–5973.

Bailey, T. L., Johnson, J., Grant, C. E., & Noble, W. S. (2015). The MEME suite. Nucleic acids research, 43(W1), W39–W49.

Barreto, P., Dambire, C., Sharma, G., Vicente, J., Osborne, R., Yassitepe, J., … & Arruda, P. (2022). Mitochondrial retrograde signaling through UCP1-mediated inhibition of the plant oxygen-sensing pathway. Current Biology, 32(6), 1403–1411.

Brestoff, J. R., Singh, K. K., Aquilano, K., Becker, L. B., Berridge, M. V., Boilard, E., … & Zheng, M. (2025). Recommendations for mitochondria transfer and transplantation nomenclature and characterization. Nature metabolism, 7(1), 53–67.

Castro-Mondragon, J. A., Riudavets-Puig, R., Rauluseviciute, I., Berhanu Lemma, R., Turchi, L., Blanc-Mathieu, R., … & Mathelier, A. (2022). JASPAR 2022: the 9th release of the open-access database of transcription factor binding profiles. Nucleic acids research, 50(D1), D165–D173.

Costa-Mattioli, M., & Walter, P. (2020). The integrated stress response: From mechanism to disease. Science, 368(6489), eaat5314.

De Clercq, I., Vermeirssen, V., Van Aken, O., Vandepoele, K., Murcha, M. W., Law, S. R., … & Van Breusegem, F. (2013). The membrane-bound NAC transcription factor ANAC013 functions in mitochondrial retrograde regulation of the oxidative stress response in Arabidopsis. The Plant Cell, 25(9), 3472–3490.

Del-Saz, N. F., Ribas-Carbo, M., McDonald, A. E., Lambers, H., Fernie, A. R., & Florez-Sarasa, I. (2018). An in vivo perspective of the role (s) of the alternative oxidase pathway. Trends in Plant Science, 23(3), 206–219.

Diaz, F. (2010). Cytochrome c oxidase deficiency: patients and animal models. Biochimica et Biophysica Acta (BBA)-Molecular Basis of Disease, 1802(1), 100–110.

El-Khoury, R., Dufour, E., Rak, M., Ramanantsoa, N., Grandchamp, N., Csaba, Z., … & Rustin, P. (2013). Alternative oxidase expression in the mouse enables bypassing cytochrome c oxidase blockade and limits mitochondrial ROS overproduction. PLoS genetics, 9(1), e1003182.

Fessler, E., Eckl, E. M., Schmitt, S., Mancilla, I. A., Meyer-Bender, M. F., Hanf, M., … & Jae, L. T. (2020). A pathway coordinated by DELE1 relays mitochondrial stress to the cytosol. Nature, 579(7799), 433–437.

Gorman, G. S., Chinnery, P. F., DiMauro, S., Hirano, M., Koga, Y., McFarland, R., … & Turnbull, D. M. (2016). Mitochondrial diseases. Nature reviews Disease primers, 2(1), 1–22.

Gray, M. W. (2012). Mitochondrial evolution. Cold Spring Harbor perspectives in biology, 4(9), a011403.

Guo, X., Aviles, G., Liu, Y., Tian, R., Unger, B. A., Lin, Y. H. T., … & Kampmann, M. (2020). Mitochondrial stress is relayed to the cytosol by an OMA1–DELE1–HRI pathway. Nature, 579(7799), 427–432.

Guo, R., Zong, S., Wu, M., Gu, J., & Yang, M. (2017). Architecture of human mitochondrial respiratory megacomplex I2III2IV2. Cell, 170(6), 1247–1257.

Gupta, S., Stamatoyannopoulos, J. A., Bailey, T. L., & Noble, W. S. (2007). Quantifying similarity between motifs. Genome biology, 8(2), R24.

Harding, H. P., Zhang, Y., Zeng, H., Novoa, I., Lu, P. D., Calfon, M., … & Ron, D. (2003). An integrated stress response regulates amino acid metabolism and resistance to oxidative stress. Molecular cell, 11(3), 619–633.

Jeong, J., Kim, J., & Kim, M. S. (2025). Dual Nature of Mitochondrial Integrated Stress Response: Molecular Switches from Protection to Pathology. Genes, 16(8), 957.

Kubat, G. B., Picone, P., Tuncay, E., Aryan, L., Girgenti, A., Palumbo, L., … & Nuzzo, D. (2025). Biotechnological approaches and therapeutic potential of mitochondria transfer and transplantation. Nature Communications, 16(1), 5709.

Kulakovskiy, I. V., Vorontsov, I. E., Yevshin, I. S., Sharipov, R. N., Fedorova, A. D., Rumynskiy, E. I., … & Makeev, V. J. (2018). HOCOMOCO: towards a complete collection of transcription factor binding models for human and mouse via large-scale ChIP-Seq analysis. Nucleic acids research, 46(D1), D252–D259.

Lageix, S., Lanet, E., Pouch-Pélissier, M. N., Espagnol, M. C., Robaglia, C., Deragon, J. M., & Pélissier, T. (2008). Arabidopsis eIF2α kinase GCN2 is essential for growth in stress conditions and is activated by wounding. BMC plant biology, 8(1), 134.

Letts, J. A., Fiedorczuk, K., & Sazanov, L. A. (2016). The architecture of respiratory supercomplexes. Nature, 537(7622), 644–648.

Lokdarshi, A., & von Arnim, A. G. (2022). Emerging roles of the signaling network of the protein kinase GCN2 in the plant stress response. Plant Science, 320, 111280.

Moore, A. L., Shiba, T., Young, L., Harada, S., Kita, K., & Ito, K. (2013). Unraveling the heater: new insights into the structure of the alternative oxidase. Annual Review of Plant Biology, 64, 637–663.

Ng, S., Ivanova, A., Duncan, O., Law, S. R., Van Aken, O., De Clercq, I., … & Giraud, E. (2013). A membrane-bound NAC transcription factor, ANAC017, mediates mitochondrial retrograde signaling in Arabidopsis. The Plant Cell, 25(9), 3450–3471.

Pakos Zebrucka, K., Koryga, I., Mnich, K., Ljujic, M., Samali, A., & Gorman, A. M. (2016). The integrated stress response. The EMBO Reports, 17(10), 1374–1395.

Quirós, P. M., Prado, M. A., Zamboni, N., D’Amico, D., Williams, R. W., Finley, D., … & Auwerx, J. (2017). Multi-omics analysis identifies ATF4 as a key regulator of the mitochondrial stress response in mammals. Journal of Cell Biology, 216(7), 2027–2045.

Roger, A. J., Muñoz-Gómez, S. A., & Kamikawa, R. (2017). The origin and diversification of mitochondria. Current Biology, 27(21), R1177–R1192.

Ryoo, H. D. (2024). The integrated stress response in metabolic adaptation. Journal of Biological Chemistry, 300(4), 107151.

Scialò, F., Fernández-Ayala, D. J., & Sanz, A. (2017). Role of mitochondrial reverse electron transport in ROS signaling: potential roles in health and disease. Frontiers in physiology, 8, 428

Selinski, J., Hartmann, A., Deckers-Hebestreit, G., Day, D. A., Whelan, J., & Scheibe, R. (2018). Alternative oxidase isoforms are differentially activated by tricarboxylic acid cycle intermediates. Plant physiology, 176(2), 1423–1432.

Stewart, J. B., & Chinnery, P. F. (2015). The dynamics of mitochondrial DNA heteroplasmy: implications for human health and disease. Nature Reviews Genetics, 16(9), 530–542.

Sweetlove, L. J., Beard, K. F., Nunes-Nesi, A., Fernie, A. R., & Ratcliffe, R. G. (2010). Not just a circle: flux modes in the plant TCA cycle. Trends in plant science, 15(8), 462–470.

Szibor, M., Dhandapani, P. K., Dufour, E., Holmström, K. M., Zhuang, Y., Salwig, I., … & Braun, T. (2017). Broad AOX expression in a genetically tractable mouse model does not disturb normal physiology. Disease models & mechanisms, 10(2), 163–171.

Szklarczyk, D., Kirsch, R., Koutrouli, M., Nastou, K., Mehryary, F., Hachilif, R., … & Von Mering, C. (2023). The STRING database in 2023: protein–protein association networks and functional enrichment analyses for any sequenced genome of interest. Nucleic acids research, 51(D1), D638–D646.

Van Aken, O., & Pogson, B. J. (2017). Convergence of mitochondrial and chloroplastic ANAC017/PAP-dependent retrograde signalling pathways and suppression of programmed cell death. Cell Death & Differentiation, 24(6), 955–960.

Vanlerberghe, G. C. (2013). Alternative oxidase: a mitochondrial respiratory pathway to maintain metabolic and signaling homeostasis during abiotic and biotic stress in plants. International journal of molecular sciences, 14(4), 6805–6847.

Vanlerberghe, G. C., & McIntosh, L. (1997). Alternative oxidase: from gene to function. Annual Review of Plant Biology, 48(1), 703–734.

Viscomi, C., & Zeviani, M. (2017). MtDNA-maintenance defects: syndromes and genes. Journal of inherited metabolic disease, 40(4), 587–599.

Viscomi, C., & Zeviani, M. (2020). Strategies for fighting mitochondrial diseases. Journal of internal medicine, 287(6), 665–684.

Wallace, D. C. (2018). Mitochondrial genetic medicine. Nature genetics, 50(12), 1642–1649.

Wang, X., & Zhang, G. (2025). The mitochondrial integrated stress response: A novel approach to anti-aging and pro-longevity. Ageing Research Reviews, 103, 102603.

Xu, Y. Z., Arrieta-Montiel, M. P., Virdi, K. S., de Paula, W. B., Widhalm, J. R., Basset, G. J., … & Mackenzie, S. A. (2011). MutS HOMOLOG1 is a nucleoid protein that alters mitochondrial and plastid properties and plant response to high light. The Plant Cell, 23(9), 3428–3441.

